# Efficiently finding activity cliffs

**DOI:** 10.1101/2025.09.17.676791

**Authors:** Akash Surendran, Ramón Alain Miranda-Quintana

**Affiliations:** Department of Chemistry and Quantum Theory Project, University of Florida, FL, Gainesville, 32611

**Keywords:** activity cliff, clustering, similarity, structure-activity relationship, chemical space

## Abstract

Activity cliffs remain a key challenge in computational drug design, defying even modern machine learning approaches. Here, we present a dual approach that allows either navigating a structure-activity landscape to quickly identify activity cliffs, or how avoid them and identify maximally smooth sectors of chemical space. Both methods are built upon the BitBIRCH clustering algorithm, so they are ideally suited to analyze very large compound libraries. The code is freely available at https://github.com/mqcomplab/BitBIRCH_AC.

## 1. INTRODUCTION

The similarity property principle is a central concept in structure-activity-relationships (SAR) and several approaches towards machine learning (ML) based drug design, asserting that structurally similar compounds tend to exhibit similar biological properties^1–4^. With the ever-increasing use of machine learning in computational and medicinal chemistry, there has been a growing interest in the topic of activity cliffs (AC), which represent abrupt discontinuities in the structure-activity landscape^5–8^. Defined as pairs or groups of structurally similar compounds that show unexpectedly pronounced differences in biological activity or potency against the same target, their dual character in medicinal chemistry^9^ in striking: while they complicate the predictive accuracy of computational methods such as ML or SAR based modelling (sometimes considered as troublesome outliers in Quantitative SAR (QSAR) studies), often leading to erroneous predictions^10^, they simultaneously encode high information content for medicinal chemists in guiding compound optimization, leading to dedicated efforts in their identification and exploitation^9,11^.

ML models, which have propelled the rational design and screening of novel compounds, are particularly challenged by the presence of ACs. Conventional algorithms frequently fail to capture these underlying discontinuities, resulting in poor predictive performance for these exceptional compound pairs. Given the activities and similarities of each molecule in the dataset, this presents an O(*N*^2^) problem to “hunt” every pair of ACs, which would make this pairwise approach intractable for large datasets. Nevertheless, this presents an intricate yet interesting challenge in ML based drug design to improve the capacity of models to identify and learn these edge cases to build more reliable computational tools for virtual screening and SAR analysis.

Recently, our group developed BitBIRCH^12,13^, a highly efficient and scalable clustering algorithm designed for handling ultra-large molecular libraries in cheminformatics and drug discovery. BitBIRCH leverages a convenient tree structure^14^ and the instant similarity (iSIM)^15,16^ formalism to use binary molecular fingerprints and Tanimoto similarity for clustering molecular datasets at unprecedented speed and scale. Here, we leverage BitBIRCH in two different ways as it pertains to the problem of ACs, showing how one could use this algorithm to either efficiently identify the presence of ACs in the data, quickly singling them out, or as a way to obtain maximally smooth sectors of chemical space, by forming clusters that are guaranteed to *not* have property discontinuities. These novel adaptations of BitBIRCH efficiently identify AC pairs, circumventing the prohibitive O(*N*^2^) complexity of naïve enumeration while preserving a comparable accuracy to pairwise analysis. This approach sets the stage for systematic and scalable identification of activity cliffs, which could aid in future generation of larger AC data for training and advancements in SAR analysis and machine learning applications in drug discovery.

## 2. METHODOLOGY

Every single definition of ACs starts by identifying pairs of molecules that are very similar. However, doing this without any preprocessing over a full library can be prohibitive due to the O(*N*^2^) of having to explore every possible pair. Given that with BitBIRCH we can group molecules much more efficiently than this, we propose a simple solution to this problem: perform BitBIRCH over the whole set, and do an exhaustive pairwise analysis only within each cluster. This effectively turns the problem of AC finding into a local search over just a few (much smaller) subsets. This greatly reduces the computational burden in multiple ways. First, because ACs are usually only of interest when the Tanimoto fingerprint similarity is greater or equal than 0.9, we will only get a handful of clusters with more than one molecule, and since we can ignore singletons, this decreases the volume of data to be analyzed. Moreover, the analysis of each cluster could be carried out in parallel, so the overall cost of this recipe is 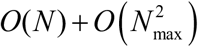 number of molecules, and *N_max_* is the number of molecules in the most populated cluster. Note that this represents a much improved scaling, especially since *N* >> *N_max_*. We can do this in a single-shot manner, but this methodology is so efficient that we can even afford to spend a bit more of time to refine the final results, in order to guarantee maximum agreement with the full pairwise method. For instance, we could run the BitBIRCH + local pairwise analysis a couple of times, removing after each iteration the pairs of molecules identified as ACs. This iterative scheme can be repeated until no new ACs are found which, in our experience, is only 2-3 rounds of BitBIRCH for a given library. If even more precision is needed, we could perform the BitBIRCH clustering using a similarity threshold decreased by some offset, in order to explore bigger clusters. That’s it, if one is interested in finding ACs at a 0.9 Tanimoto similarity, doing the clustering with a 0.8 or 0.7 threshold produces more flexible clusters, over which we could then perform a detailed pairwise analysis at the 0.9 level. In short, one could have an iterative or non-iterative analysis with or without an offset (all of these possibilities are considered in the next section).

The strategies described above are focused on finding ACs, and do not require any major modifications of the standard BitBIRCH. But what if one wants to avoid ACs and instead find maximally smooth subsets in chemical space? This can be useful to train ML models, or in property-aware contrastive learning approaches. Here, we will follow a definition inspired by that of Grisoni et al.^10^, identifying as ACs those pairs of compounds with a Tanimoto above a given threshold, and which properties differ in at least an order of magnitude. (In the following, for simplicity, we use a logarithmic scale for properties, so “at least an order of magnitude” can be rephrased as “at least a unit of difference in the log scale”.) Fortunately, learning how to avoid this behavior through a “smooth BitBIRCH” only requires two simple modifications:

1. Bit Feature (BF) structure: Besides the number of molecules in the cluster, *N_j_*, the indices of the molecules, mols_*j*_, the (linear) sum of the columns of the fingerprints cluster, **ls**_*j*_, and the cluster centroid, **c**_*j*_, we now also need to store two numbers: *p_min_* and *p_max_*, corresponding to the minimum and maximum value, respectively, of the property values of the molecules in the cluster (for convenience, stored in logarithmic scale).
2. Merge criterion: For a new cluster to be formed, now two conditions need to be fulfilled. The average similarity of the molecules cannot be less than the similarity threshold. B) |*p_max_* - *p_min_*| must be less than one.

The 1^st^ change makes sense in that now we are interested in evaluating not only structural information, but also activities, so we need to keep track of them in the BF. Then, the 2^nd^ condition guarantees that there is not a pair of molecules in the cluster whose activities differ in more than one order of magnitude. Note that we can ensure this without having to store all the property values, or checking all pairwise property differences. If the biggest possible property difference in the set (e.g., |*p_max_* - *p_min_*|) is enforced to be < 1 then, by construction, every other difference in property values will also be < 1.

To test the application of BitBIRCH to the AC problem, we used 30 CHEMBL datasets curated from the MoleculeACE platform (https://github.com/molML/MoleculeACE/tree/main), each corresponding to a different macromolecular target encompassing a diverse range of chemical and biological activities. To check the consistency of our methods across different molecular representations, we generated three different binary fingerprints: ECFP, MACCS and RDKIT. The entire collection contains about 35,000 molecules and was previously used by Grisoni et al. to benchmark ML models for AC prediction^10^, with each library having different prevalence of ACs.

## 3. RESULTS

First, we checked BitBIRCH’s ability to find ACs in different conditions. Figures 1A and 1B present a summary of the performance of the different BitBIRCH versions, including the recursive, non-recursive, and offset variants across similarity thresholds from 0.9 to 0.99. Reassuringly, all methods display a general increasing trend in accuracy ratio with similarity thresholds, almost converging to unity at high similarity thresholds. Notice also how for a given offset, the difference in retrieval rate between the recursive and non-recursive versions constantly narrows with increasing threshold across all three fingerprints. As expected, the iterative approach with an offset (0.3 in this case) exhibited higher accuracy and robustness across the entire range of thresholds and fingerprint types, with successful retrieval rates close to 100% in all cases. Figure 1C contains a simplified representation of these results, showing the ratio of each method averaged over the entire range of similarity thresholds across the three fingerprints. Note how the iterative BitBIRCH with offset is not only the most accurate but also stable to the choice of fingerprints and similarity threshold, showing only minimal deviations. It is remarkable that even the non-iterative approach proves particularly effective, recovering over 80% of ACs across all fingerprint types. The influence of offsets is particularly pronounced for MACCS and RDKIT fingerprints. Effectively, these relatively small adjustments were sufficient to push the accuracy of our methods beyond 95% across all fingerprint representations.

**Figure 1.**
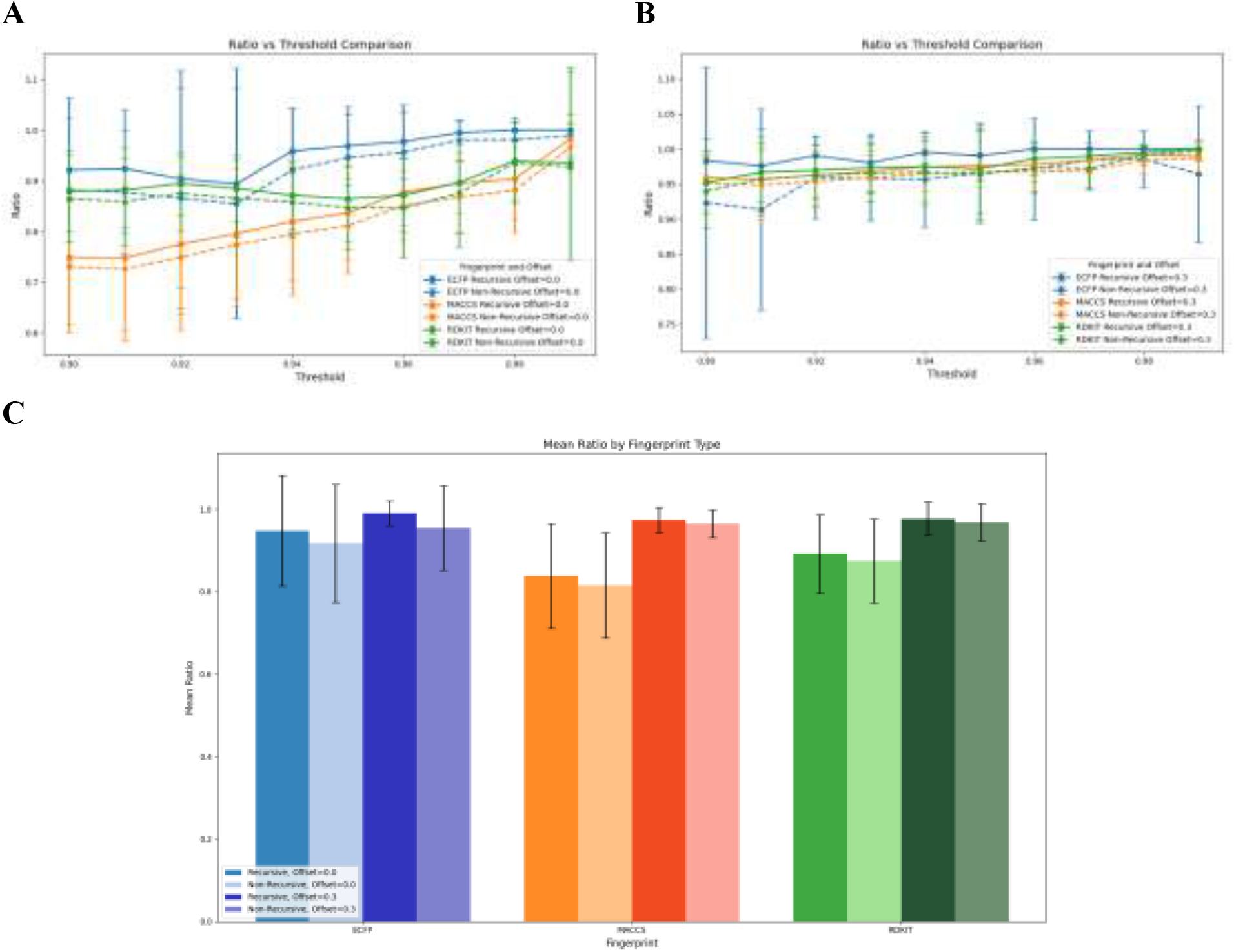
Success rate in finding ACs throughout all the libraries. A: recursive and non-recursive approaches without offset and B: recursive and non-recursive approaches with 0.3 offset, for similarity thresholds in the [0.9, 0.99] range. C: Summary of the results for all fingerprints, across similarity thresholds.

Having proved the effectiveness of BitBIRCH to find ACs, the next step is to see how we could avoid them and form clusters that have structure-activity landscapes as smooth as possible. To illustrate the changes that this introduces, we examined the top 20 most populated clusters after performing regular (diameter) BitBIRCH and smooth BitBIRCH. The results for ECFP fingerprints are shown in Figure 2A where we dissect cluster populations over different similarity thresholds. As expected, regular BitBIRCH calculations show a wide range of populations for the main clusters, with the number of molecules in the top cluster sharply decreasing with increasing similarity threshold. This is in sharp contrast with the smooth clustering results, where we can observe a relatively insensitive cluster population across similarity thresholds. Figure 2B goes into further detail about the inner structure of the different clusters. The standard deviation of the property values is much lower in the case of the smooth BitBIRCH clusters, in contrast with the regular BitBIRCH results, which can be as high as even one unit in the log scale. As proposed by Coley et al.^17,18^, the standard deviation of properties inside a cluster serves as a proxy to quantify the roughness of activity landscapes, which highlights that the new BitBIRCH variant succeeds in forming AC-free clusters.

**Figure 2.**
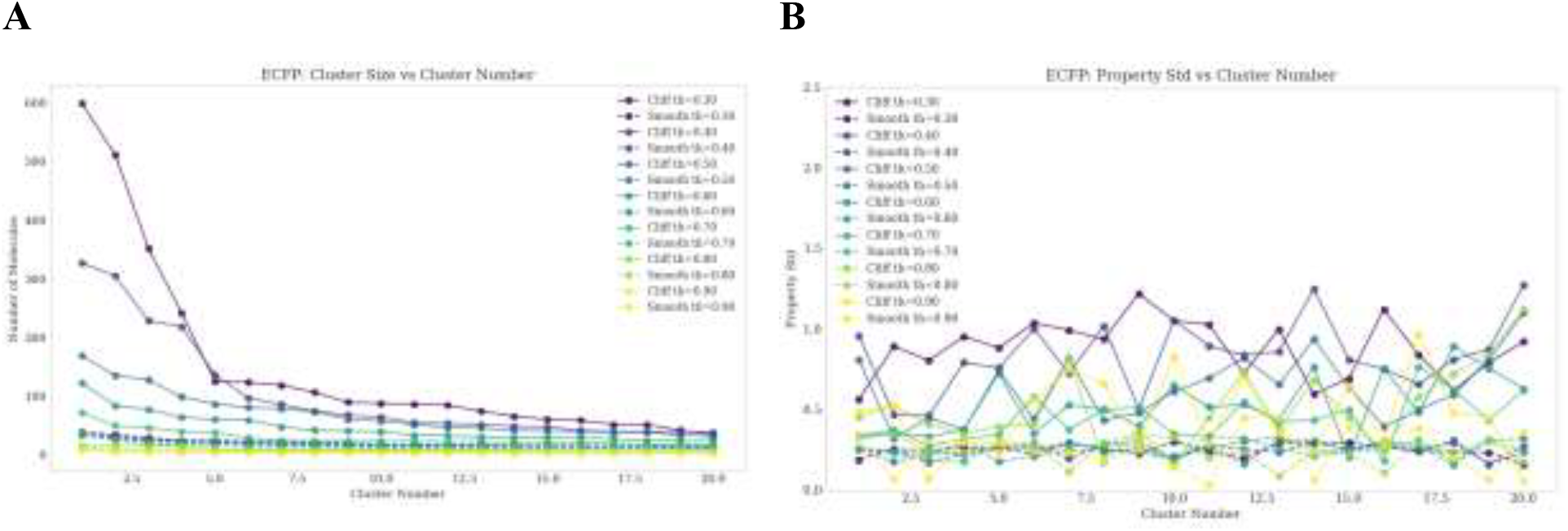
Cluster populations and standard deviations (dataset: CHEMBL234_Ki)

As a final illustration of the difference between smooth and regular BitBIRCH clusters, we compare two corresponding clusters obtained from both methods. Figure 3 shows that smooth clusters consist of molecules that share both high structural similarity and closely aligned properties, affirming their roles as regions of local SAR consistency. On the other hand, clusters containing ACs represent the edge of SAR predictability, molecules that are quite similar in their structural backbone with the presence/absence of certain functional groups inducing dramatic shifts in properties. This type of analysis is key to identify key modifications that can drive potency, while keeping a common scaffold relatively intact.

**Figure 3.**
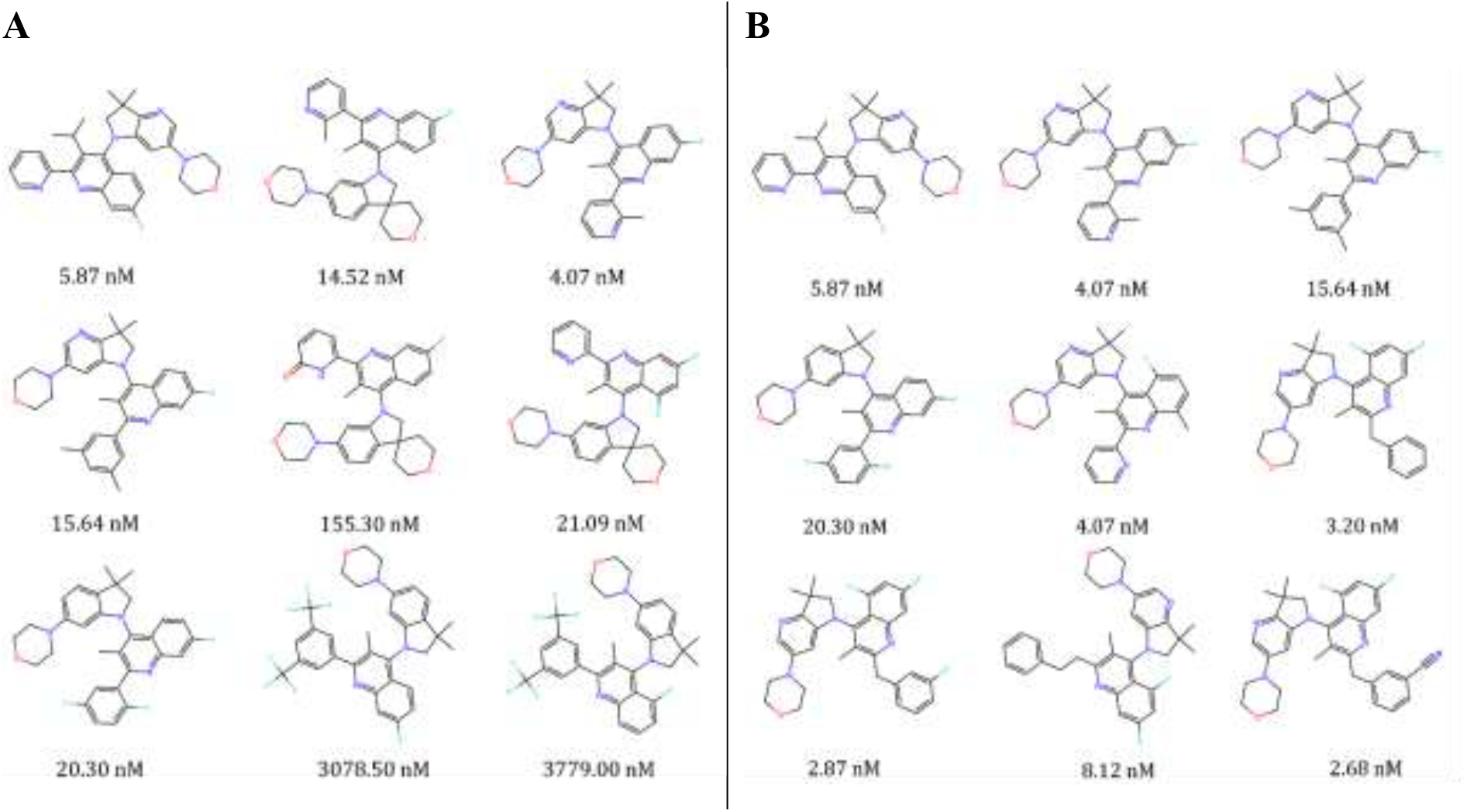
Comparison between smooth regular (AC-containing, A) and smooth (B) clusters found in the CHEMBL4005_Ki library using MACCS fingerprints.

## 4. CONCLUSIONS

We have shown how efficient clustering in fingerprint space can provide valuable information about the structure-activity relationships in a library, by quickly identifying the presence of ACs. Our BitBIRCH-powered strategies (including single-shot, non-iterative, or recursive tries, with and without similarity offsets) proved quite robust in singling out sharp discontinuities in the SAR landscape, independently of fingerprint type, and across biological targets. Moreover, to the best of our knowledge, we present the first strategy to find maximally smooth sectors in chemical space, which could be leveraged to learn better property-aware molecular representations. The direct comparison of BitBIRCH clusters with and without ACs gives clear indication of the optimal modifications that drive the potency of a given scaffold. In all of this, the key computational bottleneck is the BitBIRCH clustering, which has already been established as a hyper-efficient way to partition chemical libraries, thus making these methods particularly attractive to dissect ultra-large datasets.

## Data Availability

The AC-augmented BitBIRCH code can be foundat: https://github.com/mqcomplab/BitBIRCH_AC.

## Author Contributions

AS: Data curation, formal analysis, investigation, software, validation, visualization, and writing. RAMQ: Formal analysis, methodology, conceptualization, investigation, software, writing, funding acquisition, supervision, and resources.

## Acknowledgements

We thank support from the National Institute of General Medical Sciences and the National Institutes of Health under award number R35GM150620.

## References

(1) López-Pérez, K.; Avellaneda-Tamayo, J. F.; Chen, L.; López-López, E.; Juárez-Mercado, K. E.; Medina-Franco, J. L.; Miranda-Quintana, R. A. Molecular Similarity: Theory, Applications, and Perspectives. Artificial Intelligence Chemistry 2024, 2 (2), 100077. 10.1016/j.aichem.2024.100077.

(2) Eckert, H.; Bajorath, J. Molecular Similarity Analysis in Virtual Screening: Foundations, Limitations and Novel Approaches. Drug Discov Today 2007, 12 (5), 225–233. 10.1016/j.drudis.2007.01.011.

(3) Johnson, M. A.; Maggiora, G. M. Concepts and Applications of Molecular Similarity. (No Title) 1990.

(4) Maggiora, G.; Vogt, M.; Stumpfe, D.; Bajorath, J. Molecular Similarity in Medicinal Chemistry. J Med Chem 2014, 57 (8), 3186–3204. 10.1021/jm401411z.

(5) López-Pérez, K.; Miranda-Quintana, R. A. ICliff Taylor’s Version: Robust and Efficient Activity Cliff Determination. J Chem Inf Model 2025, 65 (11), 5801–5810. 10.1021/acs.jcim.5c00506.

(6) Stumpfe, D.; Hu, Y.; Dimova, D.; Bajorath, J. Recent Progress in Understanding Activity Cliffs and Their Utility in Medicinal Chemistry: Miniperspective. J Med Chem 2014, 57 (1), 18–28.

(7) Stumpfe, D.; Hu, H.; Bajorath, J. Evolving Concept of Activity Cliffs. ACS Omega 2019, 4 (11), 14360–14368. 10.1021/acsomega.9b02221.

(8) Dunn, T. B.; López-López, E.; Kim, T. D.; Medina-Franco, J. L.; Miranda-Quintana, R. A. Exploring Activity Landscapes with Extended Similarity: Is Tanimoto Enough? Mol Inform 2023, 42 (7), 2300056. 10.1002/minf.202300056.

(9) Cruz-Monteagudo, M.; Medina-Franco, J. L.; Pérez-Castillo, Y.; Nicolotti, O.; Cordeiro, M. N. D. S.; Borges, F. Activity Cliffs in Drug Discovery: Dr Jekyll or Mr Hyde? Drug Discov Today 2014, 19 (8), 1069–1080. 10.1016/j.drudis.2014.02.003.

(10) van Tilborg, D.; Alenicheva, A.; Grisoni, F. Exposing the Limitations of Molecular Machine Learning with Activity Cliffs. J Chem Inf Model 2022, 62 (23), 5938–5951. 10.1021/acs.jcim.2c01073.

(11) Daoud, S.; Taha, M. Protein Characteristics Substantially Influence the Propensity of Activity Cliffs among Kinase Inhibitors. Sci Rep 2024, 14 (1), 9058. 10.1038/s41598-024-59501-w.

(12) Pérez, K. L.; Jung, V.; Chen, L.; Huddleston, K.; Miranda-Quintana, R. A. BitBIRCH: Efficient Clustering of Large Molecular Libraries. Digital Discovery 2025, 4 (4), 1042– 1051.

(13) LÓpez Pérez, K.; Huddleston, K.; Jung, V.; Miranda-Quintana, R. A. BitBIRCH Clustering Refinement Strategies. J Chem Inf Model 2025.

(14) Zhang, T.; Ramakrishnan, R.; Livny, M. BIRCH: An Efficient Data Clustering Method for Very Large Databases. ACM sigmod record 1996, 25 (2), 103–114.

(15) López-Pérez, K.; Kim, T. D.; Miranda-Quintana, R. A. ISIM: Instant Similarity. Digital Discovery 2024, 3 (6), 1160–1171.

(16) Lopez-Perez, K.; Zhao, B.; Miranda-Quintana, R. A. ISIM-Sigma: Efficient Standard Deviation Calculation for Molecular Similarity. J Chem Inf Model 2025.

(17) Aldeghi, M.; Graff, D. E.; Frey, N.; Morrone, J. A.; Pyzer-Knapp, E. O.; Jordan, K. E.; Coley, C. W. Roughness of Molecular Property Landscapes and Its Impact on Modellability. J Chem Inf Model 2022, 62 (19), 4660–4671. 10.1021/acs.jcim.2c00903.

(18) Graff, D. E.; Pyzer-Knapp, E. O.; Jordan, K. E.; Shakhnovich, E. I.; Coley, C. W. Evaluating the Roughness of Structure–Property Relationships Using Pretrained Molecular Representations. Digital Discovery 2023, 2 (5), 1452–1460. 10.1039/D3DD00088E.

