## Supporting Information for "Efficiently finding activity cliffs"

**S1. Effect of offset on the performance of BitBIRCH AC**

We systematically investigated how varying offsets could influence the efficiency of AC detection across all 30 datasets and the three fingerprint types, using color gradient to represent the fraction of activity cliffs detected in Figure S1. As expected, regions corresponding to high threshold and offset display near-complete accuracy, reflecting an expanded searchable chemical space with increasing flexibility. Notably, the MACCS fingerprint shows the clearest distribution gradient among all representations, consistently increasing in accuracy as we move from lower to higher threshold and offset values. Table S1 summarizes the time taken to perform AC calculations at a given offset, including the exhaustive pairwise calculations used as the reference for estimating the success ratio (we can assume the exhaustive pairwise comparison time to be constant since the number of pairs compared is independent of threshold or offset). An offset of 0.3 takes just 14s more than the zero-offset baseline, demonstrating robust performance across fingerprint types and practical time constraints. These results underscore the importance of optimizing the offset parameter to maximize successful AC detection across diverse chemical datasets and molecular representations, while maintaining computational efficiency.

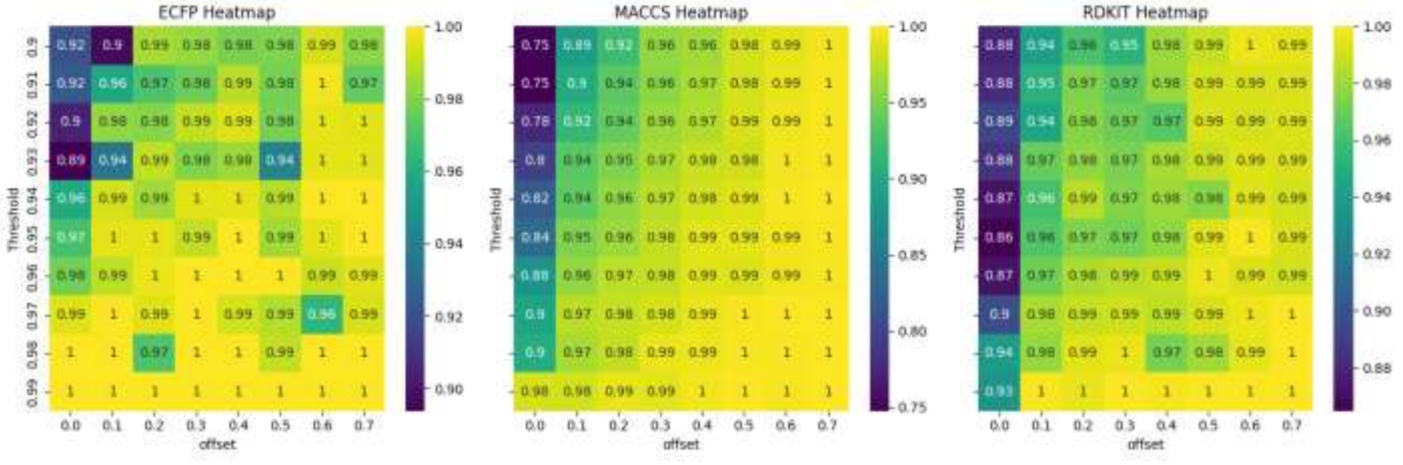

**Figure S1:** Heatmap showing the success ratio in finding ACs across different thresholds and offsets, averaged over all 30 CHEMBL datasets.

| Offset | Time(s) |
|---|---|
| 0.0 | 192 |
| 0.1 | 185 |
| 0.2 | 190 |
| 0.3 | 205 |
| 0.4 | 263 |
| 0.5 | 382 |
| 0.6 | 545 |
| 0.7 | 974 |

**Table S1:** Time taken by the iterative version of the algorithm to enumerate activity cliffs across all 30 CHEMBL libraries and three fingerprints for a fixed offset.

### S2. Cluster population and Property standard deviation across fingerprint types

Figure S2 demonstrates the cluster population and property standard deviations of the top 20 most populated smooth and regular diameter BitBIRCH clusters. Smooth clusters exhibit a more uniform and substantially lower molecule count across clusters, whereas regular diameter BitBIRCH clusters show a pronounced population imbalance: the first cluster contains a disproportionately higher number of molecules, followed by a rapid decrease in cluster population. Interestingly, only a single cluster was formed for the MACCS fingerprint, indicating that all molecules in this dataset surpass the chosen similarity threshold of 0.3 in this representation.

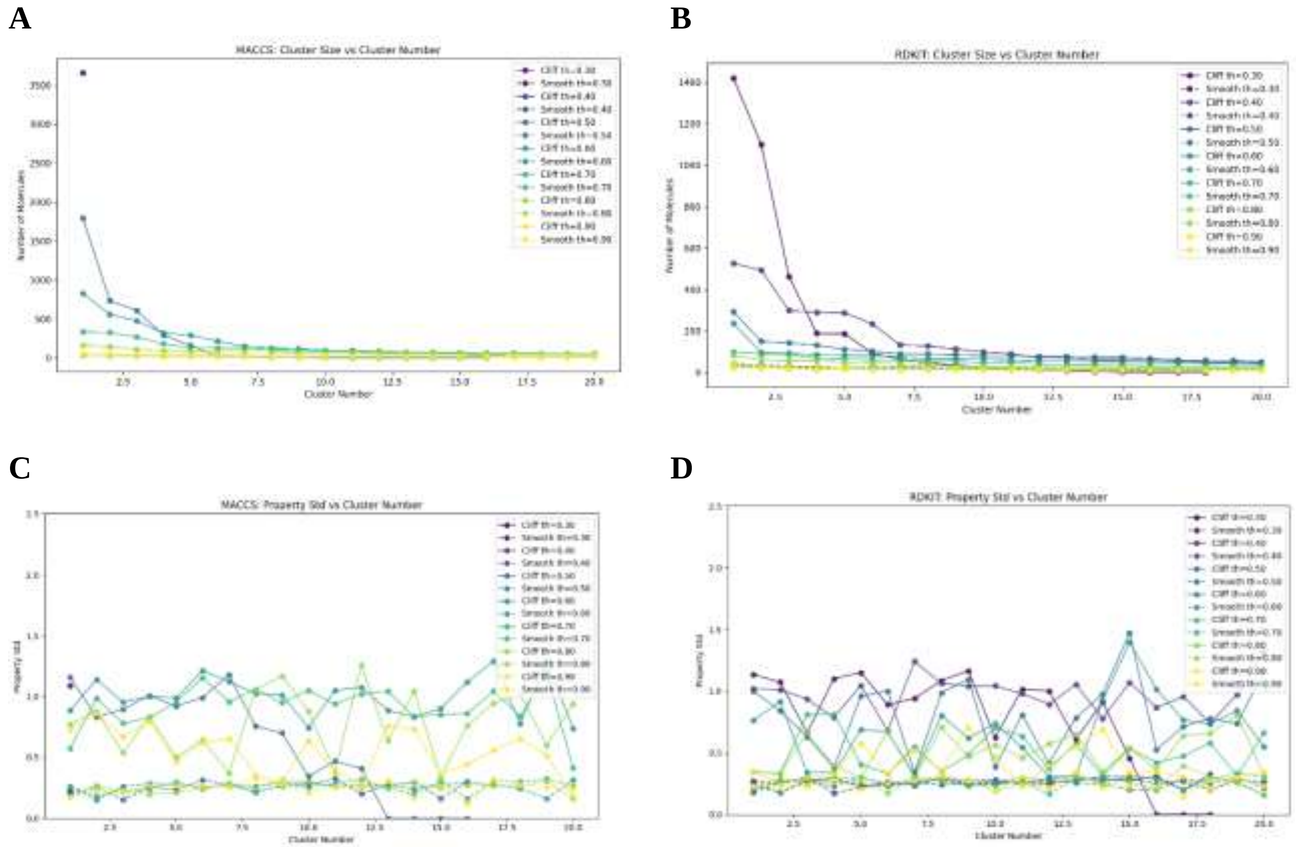


**Figure S2:** Cluster population (A and B) and property standard deviation (C and D) of smooth and regular diameter BitBIRCH clusters for the dataset CHEMBL214_Ki across MACCS and RDKIT fingerprints.

As highlighted in the main text, property fluctuations (measured as standard deviations in log units) are significantly smaller in the smooth clusters due to the absence of activity cliffs. A key observation is that the standard deviations generally decrease with increasing with increasing similarity threshold illustrating that higher structural similarity correlates with greater property similarity or the molecular similarity principle. However, some clusters even display standard deviations greater than 1, underscoring the presence and impact of activity cliffs.
